# Freezing Diluted Bovine Serum Albumin Standards Does Not Significantly Affect Standard Curves

**DOI:** 10.1101/2024.06.26.600904

**Authors:** Rachelle Sheets, Bhavik Rajaboina, Caitlin E. Bromberg, Liam P. Curtin, Mitchell L. Haddock, Phillip Stafford, Theresa Currier Thomas, Adrienne C. Scheck

## Abstract

Total protein isolation followed by quantitation is a common protocol in many laboratories. Quantitation is often done using a colorimetric assay such as the bicinchoninic acid (BCA) assay in which a change in the color of the BCA reagent is related to protein concentration. Extracted protein samples are compared to a standard curve made with dilutions of a protein standard such as bovine serum albumin (BSA) to determine their concentrations. A series of experiments was designed to determine the most reproducible and accurate method for quantifying protein concentrations of samples in an experimental series over time. The effect of freezing on diluted standards was investigated. Standards were frozen at −20°C or −80°C and serially thawed and refrozen up to three times prior to their use in a BCA assay. Thawing and refreezing the standards had no significant effect on protein concentration and the resulting standard curves. Inter-person and intra-person variability in the preparation of standards was also investigated. Protein concentration differences due to inter-person and intra-person variability were greater than protein concentration variability resulting from freezing and thawing, regardless of the freezing temperature. The most reproducible and accurate method for determining the protein concentration of extracted samples in an experimental series over time is diluting a large batch of BSA standards and freezing them at either −20°C or −80°C. Reproducibility was maintained with up to three freeze-thaws.

**Highlights:**

- Freezing diluted BSA standards at either −20°C or −80°C and thawing and refreezing them up to three times does not significantly alter their protein concentrations.
- Inter-person variability in standard curves is greater than intra-person variability, and controlling for developer decreases variability in most cases.
- There is more consistency in standard curves when a single batch of diluted standards is aliquoted and frozen at either −20°C or −80°C and thawed up to three times than when the same investigator or different investigators make fresh standards before each assay.

## Introduction

Total protein isolation followed by quantitation is a frequently used protocol in many laboratories. When establishing a protein quantification protocol, optimal reproducibility and rigor are fundamental to ensuring accurately calculated sample protein concentrations. Quantitation is often done using a colorimetric assay such as the bicinchoninic acid (BCA) assay in which a change in the color of the BCA reagent is related to protein concentration. Sample concentrations are compared to a standard curve made with dilutions of a protein standard such as bovine serum albumin (BSA) to determine their concentrations.

One commonly used kit for protein quantitation is the Pierce™ BCA Protein Assay Kit which is available with pre-diluted BSA standards or undiluted BSA that can be diluted to the desired concentration range. Although the protocol for the assay using undiluted BSA calls for storing the undiluted BSA vials at room temperature and diluting the standards before use (Thermo Scientific™ Pierce™ BCA Protein Assay Kits Catalog numbers: 23225 and 23227), many researchers find it more efficient to aliquot batches of diluted BSA standards and freeze them for future assays, sometimes using each aliquot several times. Benefits of freezing diluted BSA standards include improved intra-person and inter-person consistency across assays performed at different times to quantify all samples in an experiment. This is particularly important when new trainees are assigned to run protein assays, increasing the risk of pipettor or dilution errors that can compromise results. While a number of publications have compared different quantitation methods, the importance of preparation and storage methods for the assay standards has generally not been addressed ^1-4^. Some publications and online guides indicate that storage conditions and freeze-thaw cycles of proteins result in degradation of protein due to denaturing through temperature-induced structural changes, protein aggregation, and changes in concentration during freeze thaw events; however, these reports typically look at complex mixtures of proteins isolated from cells or tissue and not a purified single protein such as those used as standards ^2^. Researchers who choose to freeze diluted standards typically store them at either −20°C or −80°C. Freezing at −80°C is thought to improve stability and protein preservation due to slower degradation and reduced aggregation in comparison to storing at −20°C. However, −80°C storage is expensive and space is typically limited, so some researchers choose to store these standards at −20°C. While preferences were identified online, no controlled evaluation of fresh BSA standards compared to frozen diluted BSA standards, storage temperature (−20°C or −80°C) or number of freeze-thaw events was identified in the literature.

## Methods

A series of experiments was designed to determine the most reproducible and accurate method for quantifying protein concentrations of samples in an experimental series over time. We compared results from assays performed by a single investigator versus multiple investigators using fresh standards, standards frozen at -20°C or -80°C, and multiple freeze-thaw cycles. Dependent variables included standard curves from BCA assays and sample protein concentrations. Confounding variables that were controlled in these experiments included length of time frozen, standard preparation variability, and developer variability (combining of reagents A and B to develop the plate).

The Pierce™ BCA Protein Assay Kit (CAT #: 23250, Thermo Scientific™) was used for all assays following the manufacturer’s protocol for the preparation of the BCA developer solution and incubation times. Eight dilutions of BSA standards made in double-distilled H_2_O with concentrations ranging from 0-420µg/ml were used for all standard curves. For freezing, diluted standards were aliquoted into 2 milliliter (ml) screwcap microtubes (Sarstedt, Newton, NC, product number 72.694.306) and frozen at −20° C or −80° C. When the effect of thawing and refreezing on diluted standards was investigated, standards were thawed and refrozen every 7 days. On the days the assays were run, standards and samples were thawed on ice and used immediately once thawed. All standards and samples were analyzed in triplicate. Absorbance was read at 565 nanometers (nm) on an Epoch Microplate Spectrophotometer (Agilent Technologies, Inc., Santa Clara, CA) using Gen5 Data Analysis software. Data was analyzed using GraphPad Prism version 10.2.1 for Windows (GraphPad Software, Boston, MA). All standards were plotted as an average of the data ± standard error of the mean (SEM) after outliers were removed from the data sets. A single outlier was excluded from Figure 2B (180μg/ml standard) due to an A_565_ value 4 standard deviations from the mean. Additional outliers were identified by GraphPad using the ROUT method (Q = 1%). Single outliers were found in Figure 3, investigator 1 - 60µg/ml standard; Figure 3 investigator 3 - 0µg/ml standard and Figure 6, investigator 3 - 240µg/ml standard. The 300µg/ml standard data was excluded from Figure 5, set 3 due to a technical error. Standard curves were made using second order polynomial trendlines resulting in R^2^ values ≥0.98. Protein concentrations were extrapolated from this curve. The coefficient of variation (CV) for the protein concentrations was determined for each standard preparation condition using Microsoft Excel (Microsoft, Redmond, WA). Typically, a CV ≤ 10% is considered to be in the upper 99^th^ percentile of performance ^5^.

**Figure 1.**
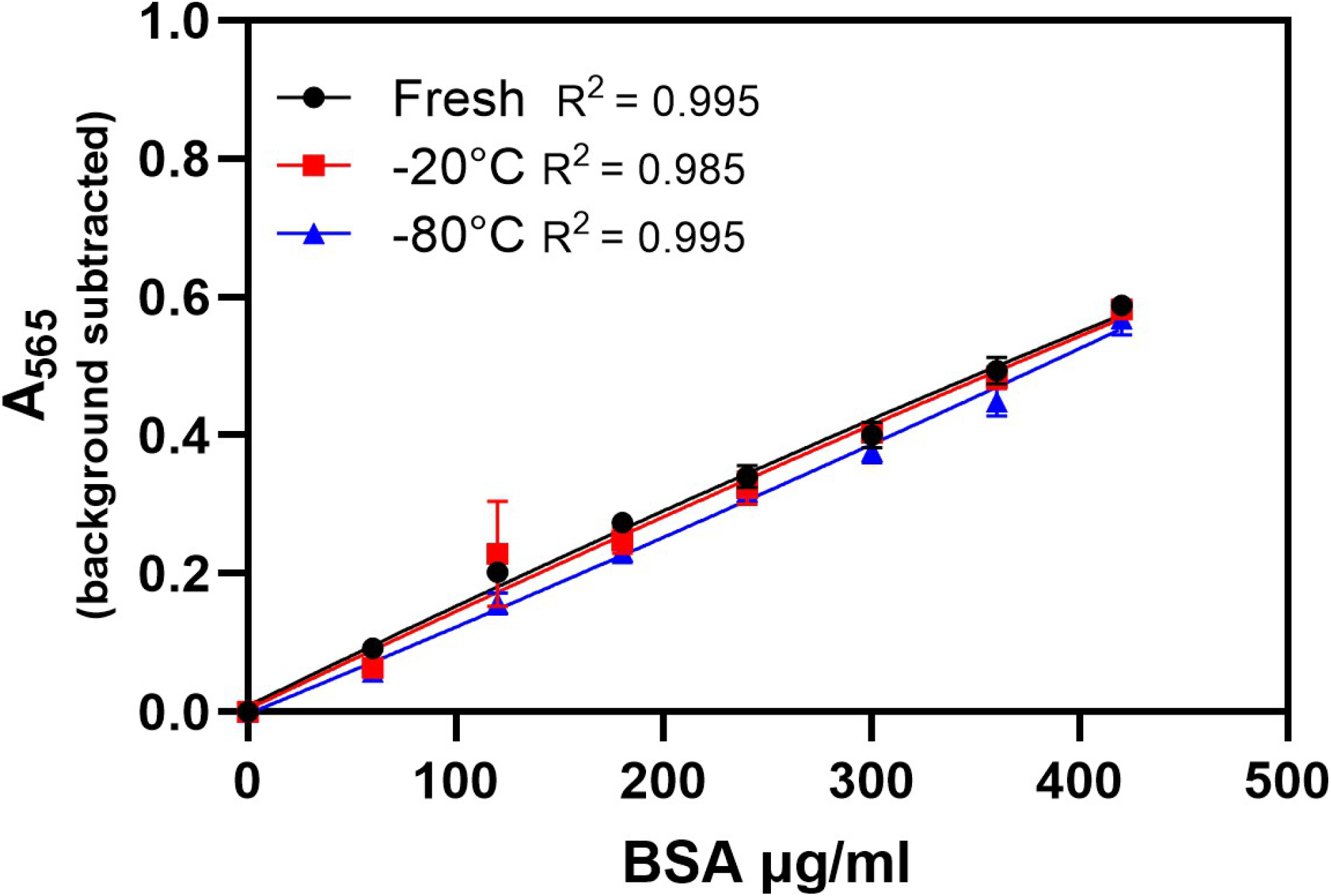
Standards prepared by one investigator were used fresh or following a single freeze at -20°C or -80°C to create standard curves. The CV of the sample protein concentrations using the freshly made and frozen standards was low (0.02 and 0.04). There was no statistically significant difference in the interpolated protein concentrations of samples when the fresh and previously frozen standards were used to make the standard curve. Data is mean ± SEM.

**Figure 2.**
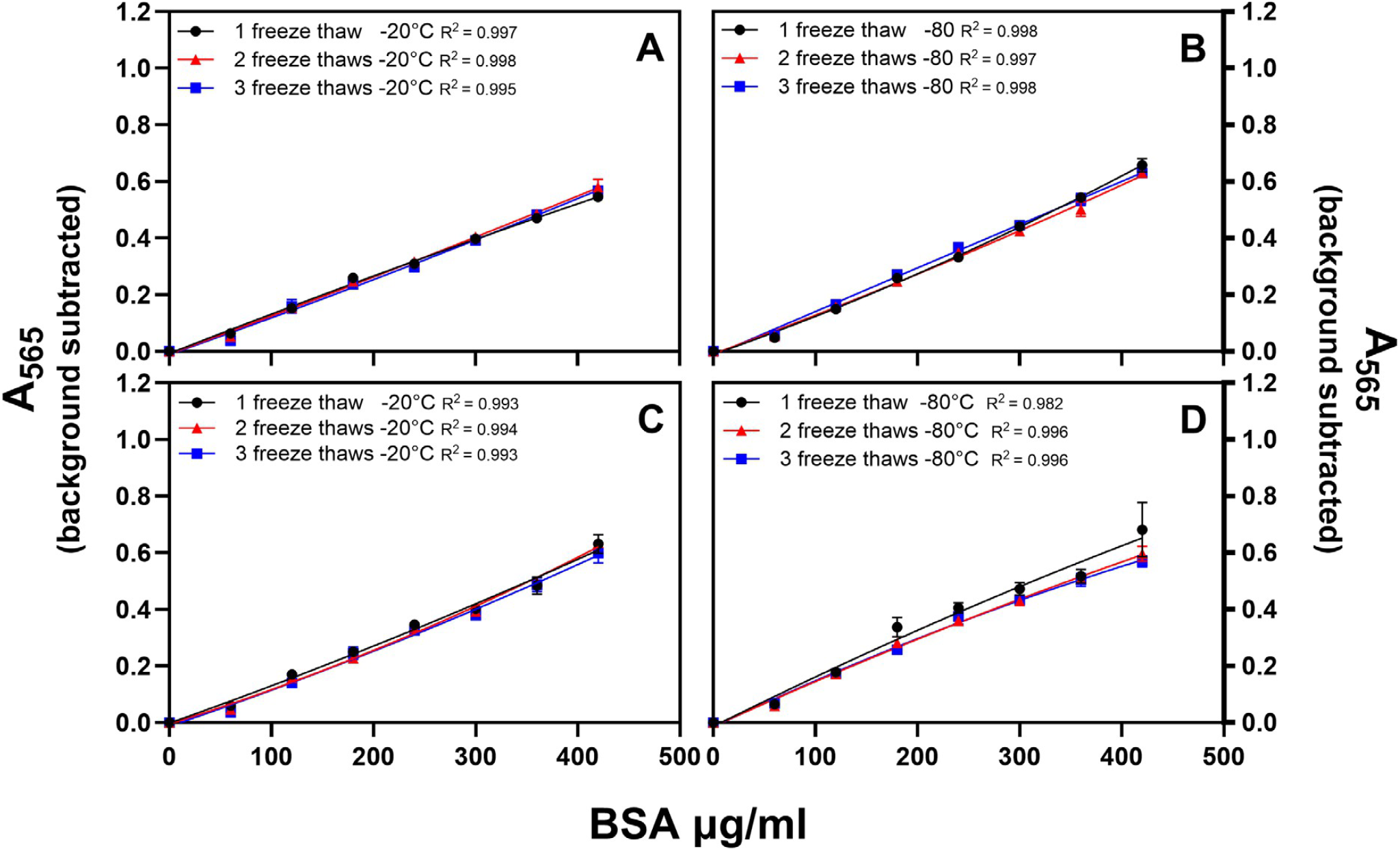
Diluted standards prepared by one investigator were frozen at -20°C or -80°C, then thawed and refrozen every 7 days up to three times. (A, B) Standards frozen at -20°C were assayed 3 days before standards frozen at -80°C. (C, D) Standards frozen at -20°C and -80°C were assayed on the same day. There was no statistically significant difference in the interpolated protein concentrations of the hypothetical samples when each set of standards was used to make the standard curve. Data is mean ± SEM.

**Figure 3.**
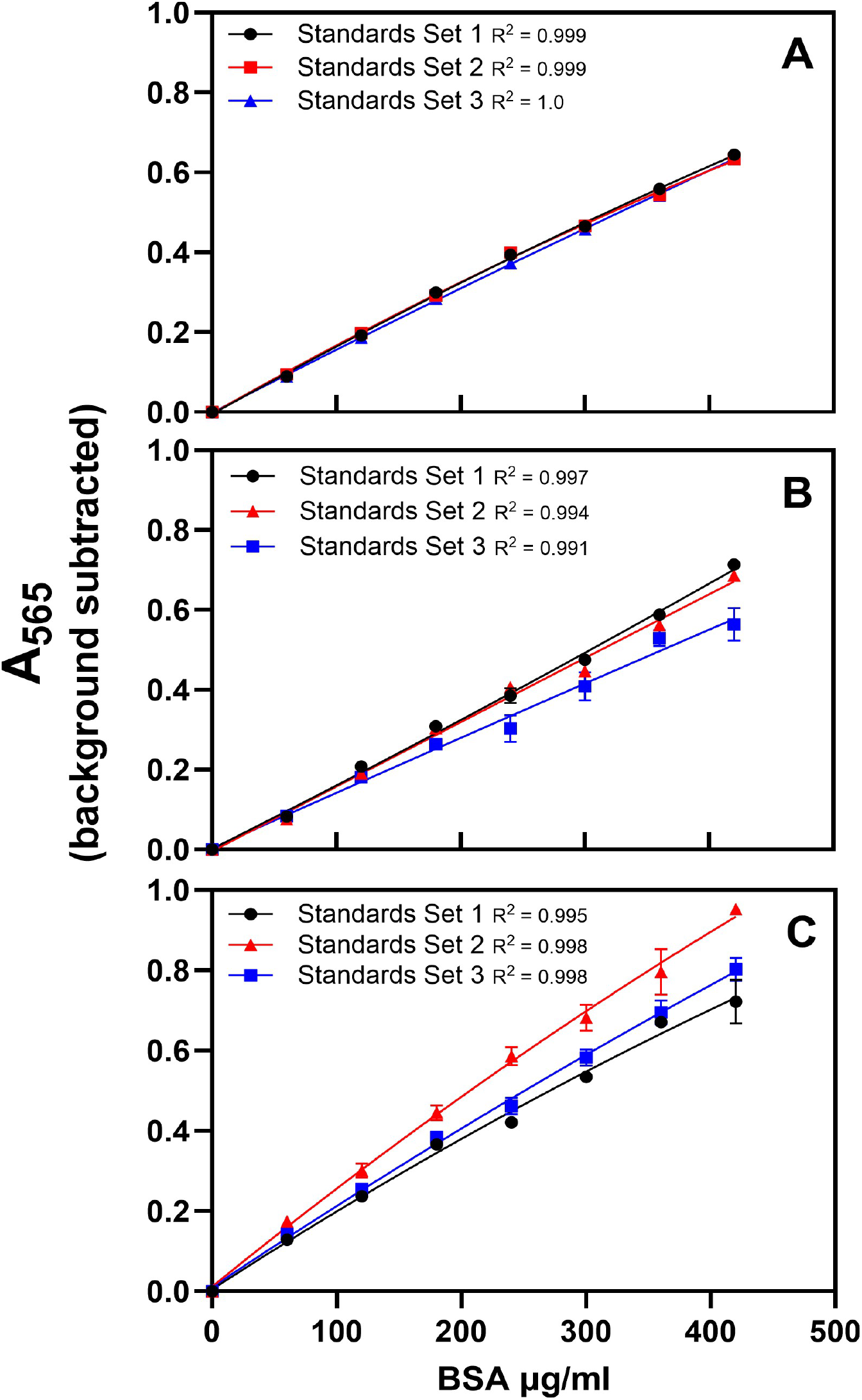
Standard curves made by three experienced investigators on 3 separate occasions were compared to assess intra-person variability (panels A, B, and C). Each investigator prepared three curves, and standards were plated in triplicate. (A) The slopes of the curves were identical for investigator 1, yielding matching standard curves and a low CV (0.01-0.02) when the highest and lowest hypothetical samples at the edges of the curve are excluded. (B) Two of the three slopes were identical for investigator 2, with CV ranging from 0.06-0.10 when the highest and lowest hypothetical samples at the edges of the curve are excluded. (C) Data from investigator 3 showed the most intra-person variability with CV values ranging from 0.13-0.15 irrespective of the values from the highest and lowest hypothetical samples. Data is mean ± SEM.

**Figure 4.**
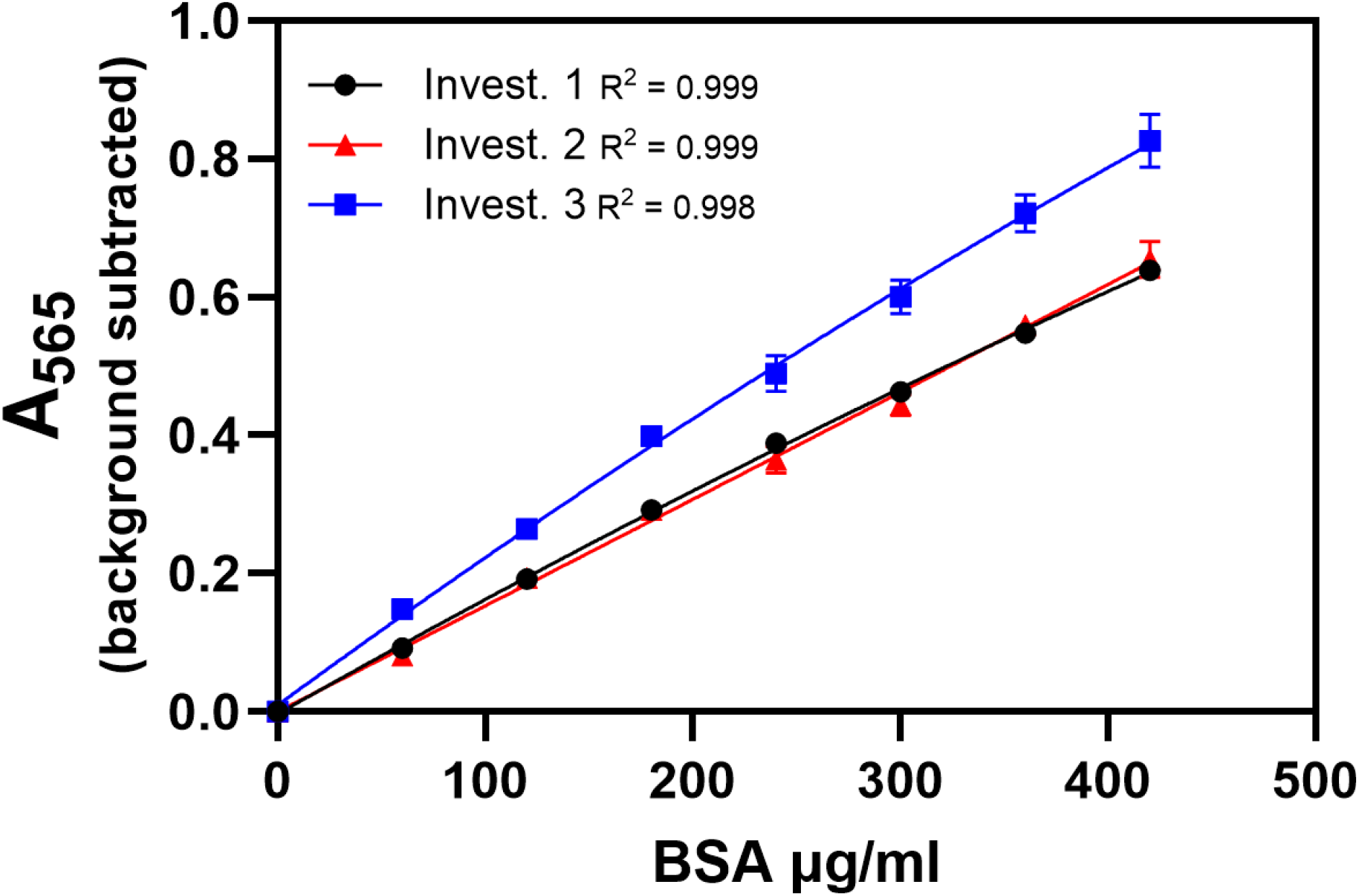
The average of the individual investigators’ data (Figure 3) was used to create a single standard curve from each investigator to assess inter-person variability. CV values ranged from 0.09-0.12 when the 0.1 hypothetical values were omitted. Data is mean ± SEM.

**Figure 5.**
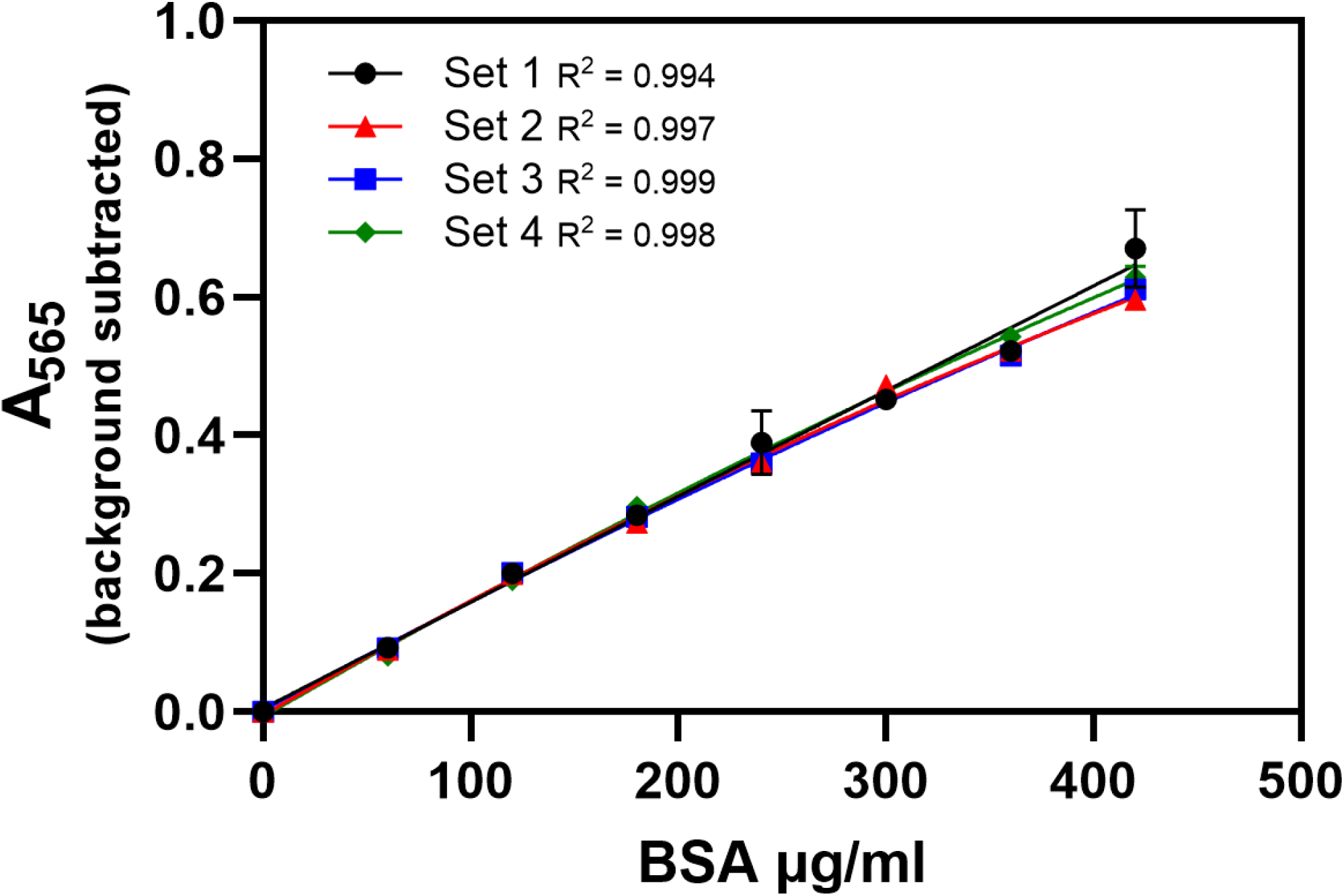
Four sets of diluted standards were prepared by one investigator and assayed on the same plate to assess intra-person variability when plating and developer were controlled. When developer variability was controlled in this way, the curves for each set of standards prepared by one investigator were aligned for protein concentrations ≤240µg/ml. Data is mean ± SEM.

**Figure 6.**
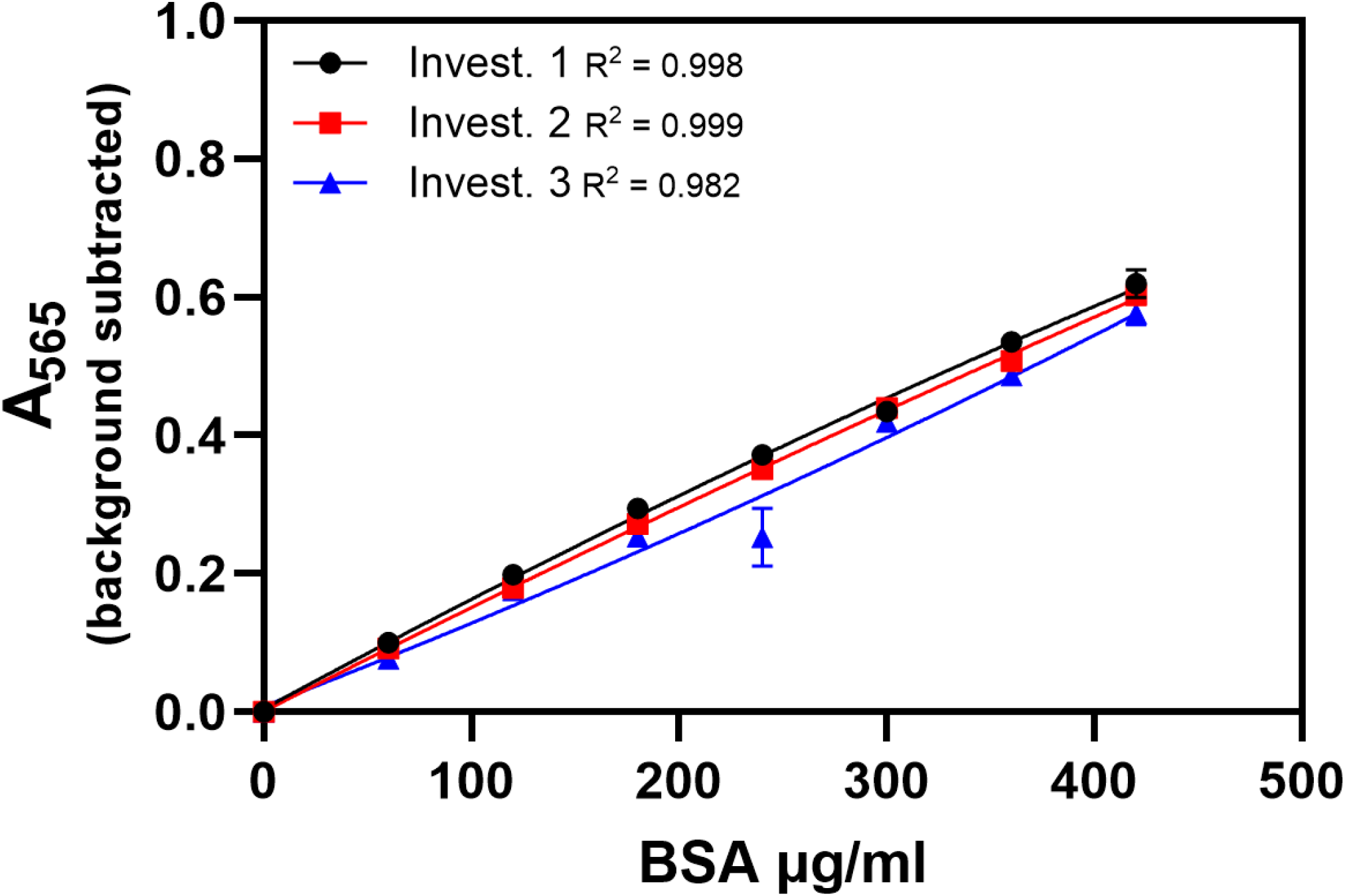
Standard curves made by three investigators, plated by one person, and assayed on the same plate were compared to assess inter-person variability while plating and developer variability were controlled. Each investigator made one curve. Inter-person variability in interpolated protein concentrations was greater than intra-person variability (Figure 5, Table 5), although there was not a statistically significant difference between Investigators1 and 2. Data is mean ± SEM.

## Results

### Effect of Freezing on Diluted BSA Standards

One investigator diluted, aliquoted, and froze standards. A fresh set of standards was made by the same investigator when the frozen standards were thawed (1^st^ thaw). The calculated protein concentration of two samples was determined using the frozen and freshly diluted standards. To ensure that the calculated protein concentration would be within the range of the assay, the samples were diluted 1:20. To evaluate reproducibility of sample dilution, two 1:20 dilutions were made for each unknown sample, and each was assayed in triplicate. The average absorbance of the two dilutions was used to determine calculated protein. The CV of the protein concentrations of the samples using the freshly made and frozen standards were low (0.02 and 0.04). There was no statistically significant difference in the interpolated protein concentrations of samples when the fresh and previously frozen standards were used to make the standard curve. This was true whether standards were frozen at −20°C or −80°C (Figure 1, Table 1).

**Table 1.**
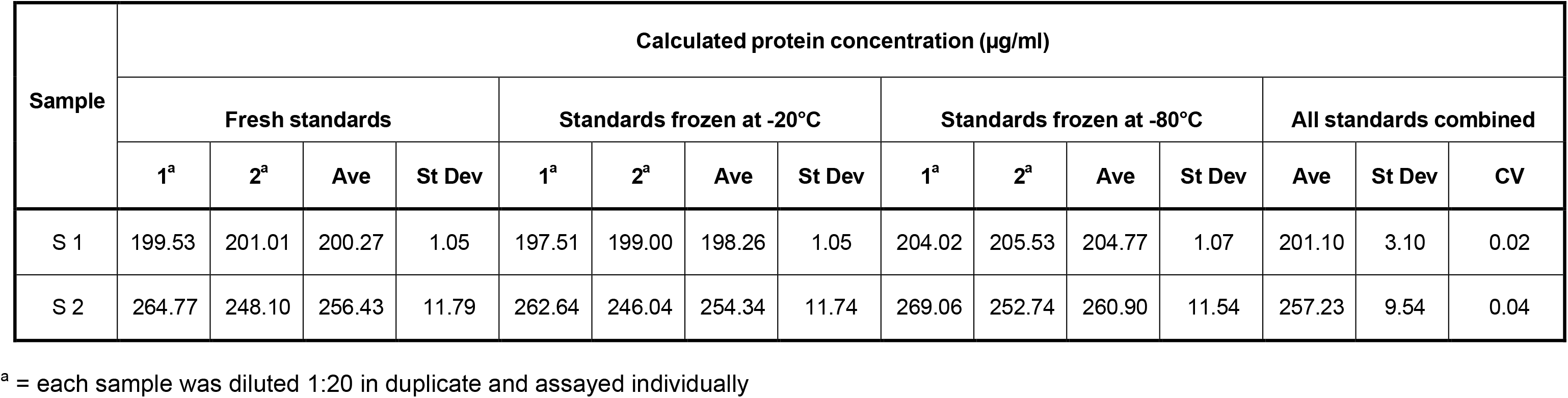
Standards prepared by a single investigator were used fresh or following a single freeze at -20°C or -80°C to create standard curves as described in the methods. Two samples (S1, S2) were diluted 1:20 in duplicate, each dilution was assayed in triplicate and the calculated protein concentration was determined using the frozen and freshly diluted standards.

| Sample | Calculated protein concentration (µg/ml) |  |  |  |  |  |  |  |  |  |  |  |  |  |  |
| --- | --- | --- | --- | --- | --- | --- | --- | --- | --- | --- | --- | --- | --- | --- | --- |
|  | Fresh standards |  |  |  | Standards frozen at -20°C |  |  |  | Standards frozen at -80°C |  |  |  | All standards combined |  |  |
|  | 1<sup>a</sup> | 2<sup>a</sup> | Ave | St Dev | 1<sup>a</sup> | 2<sup>a</sup> | Ave | St Dev | 1<sup>a</sup> | 2<sup>a</sup> | Ave | St Dev | Ave | St Dev | CV |
| S 1 | 199.53 | 201.01 | 200.27 | 1.05 | 197.51 | 199.00 | 198.26 | 1.05 | 204.02 | 205.53 | 204.77 | 1.07 | 201.10 | 3.10 | 0.02 |
| S 2 | 264.77 | 248.10 | 256.43 | 11.79 | 262.64 | 246.04 | 254.34 | 11.74 | 269.06 | 252.74 | 260.90 | 11.54 | 257.23 | 9.54 | 0.04 |

<sup>a</sup> = each sample was diluted 1:20 in duplicate and assayed individually

### Effect of Thawing and Refreezing on Diluted BSA Standards

Standards frozen at either −20°C or −80°C and thawed once, twice, or three times were used to create standard curves. Standards frozen at −80°C were assayed three days after the standards frozen at −20°C using reagents from the same kit to develop the plates (Figures 2 A and B, Tables 2 A and B). Eight absorbance values (0.1 to 0.8) were used as hypothetical samples. This experiment was repeated with standards frozen at −80°C and −20°C assayed on the same day using one batch of developer to control for both time frozen and developer variability (Figures 2 C and D, Table 2 C). There was no significant difference in standard curves between the two experiments. With the exception of the 0.1 hypothetical samples, the CV was low for each sample (0.01-0.1). There was no statistically significant difference in the interpolated protein concentrations of the hypothetical samples when each set of standards was used to make the standard curve.

**Table 2a and 2b.** Diluted standards prepared by one investigator were frozen at -20°C and -80°C, then thawed and refrozen every 7 days up to three times. Standards frozen at -20°C were assayed 3 days before standards frozen at -80°C. All assays were done in triplicate. Eight absorbance values (0.1 – 0.8) were used as hypothetical samples and protein concentrations were calculated using each of the thawed standards.

| Hypothetical Sample ($A_{565}$) | Calculated protein concentration ($\mu\text{g/ml}$)<br>-20°C Standards | | | | | |
| --- | --- | --- | --- | --- | --- | --- |
|  | 1 Thaw | 2 Thaws | 3 Thaws | Ave | St Dev | CV |
| 0.10 | 13.68 | 15.57 | 16.39 | 15.21 | 1.39 | 0.09 |
| 0.20 | 85.59 | 90.17 | 93.50 | 89.75 | 3.97 | 0.04 |
| 0.30 | 159.28 | 163.31 | 168.22 | 163.60 | 4.48 | 0.03 |
| 0.40 | 234.87 | 235.05 | 240.75 | 236.89 | 3.35 | 0.01 |
| 0.50 | 312.54 | 305.47 | 311.29 | 309.77 | 3.77 | 0.01 |
| 0.60 | 392.45 | 374.65 | 379.98 | 382.36 | 9.14 | 0.02 |
| 0.70 | 474.82 | 442.65 | 446.97 | 454.81 | 17.46 | 0.04 |
| 0.80 | 559.88 | 509.53 | 512.37 | 527.26 | 28.29 | 0.05 |

| Hypothetical Sample ($A_{565}$) | Calculated protein concentration ($\mu\text{g/ml}$)<br>-80°C Standards | | | | | |
| --- | --- | --- | --- | --- | --- | --- |
|  | 1 Thaw | 2 Thaws | 3 Thaws | Ave | St Dev | CV |
| 0.10 | 17.08 | 15.49 | 15.91 | 16.16 | 0.83 | 0.05 |
| 0.20 | 91.06 | 88.50 | 80.41 | 86.65 | 5.56 | 0.06 |
| 0.30 | 159.50 | 158.52 | 145.16 | 154.39 | 8.01 | 0.05 |
| 0.40 | 223.50 | 225.90 | 210.18 | 219.86 | 8.47 | 0.04 |
| 0.50 | 283.82 | 290.90 | 275.45 | 283.39 | 7.73 | 0.03 |
| 0.60 | 341.03 | 353.78 | 340.99 | 345.27 | 7.37 | 0.02 |
| 0.70 | 395.56 | 414.71 | 406.81 | 405.69 | 9.62 | 0.02 |
| 0.80 | 447.77 | 473.88 | 472.89 | 464.85 | 14.80 | 0.03 |

**Table 2c.**
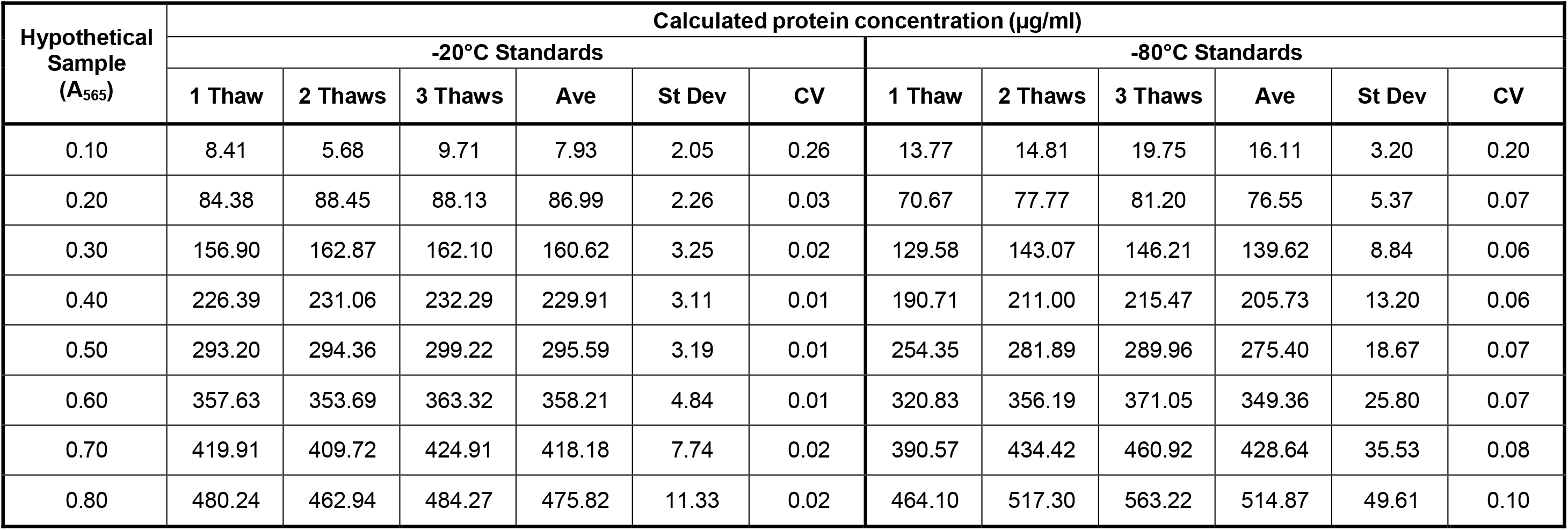
Diluted standards prepared by one investigator were frozen at -20°C and -80°C, then thawed and refrozen every 7 days up to three times. Standards frozen at -20°C and -80°C were assayed on the same day. All assays were done in triplicate. Eight absorbance values (0.1 – 0.8) were used as hypothetical samples and protein concentrations were calculated using each of the thawed standards.

| Hypothetical<br>Sample<br>(A<sub>565</sub>) | Calculated protein concentration (µg/ml) |  |  |  |  |  |  |  |  |  |  |  |
| --- | --- | --- | --- | --- | --- | --- | --- | --- | --- | --- | --- | --- |
|  | -20°C Standards |  |  |  |  |  | -80°C Standards |  |  |  |  |  |
|  | 1 Thaw | 2 Thaws | 3 Thaws | Ave | St Dev | CV | 1 Thaw | 2 Thaws | 3 Thaws | Ave | St Dev | CV |
| 0.10 | 8.41 | 5.68 | 9.71 | 7.93 | 2.05 | 0.26 | 13.77 | 14.81 | 19.75 | 16.11 | 3.20 | 0.20 |
| 0.20 | 84.38 | 88.45 | 88.13 | 86.99 | 2.26 | 0.03 | 70.67 | 77.77 | 81.20 | 76.55 | 5.37 | 0.07 |
| 0.30 | 156.90 | 162.87 | 162.10 | 160.62 | 3.25 | 0.02 | 129.58 | 143.07 | 146.21 | 139.62 | 8.84 | 0.06 |
| 0.40 | 226.39 | 231.06 | 232.29 | 229.91 | 3.11 | 0.01 | 190.71 | 211.00 | 215.47 | 205.73 | 13.20 | 0.06 |
| 0.50 | 293.20 | 294.36 | 299.22 | 295.59 | 3.19 | 0.01 | 254.35 | 281.89 | 289.96 | 275.40 | 18.67 | 0.07 |
| 0.60 | 357.63 | 353.69 | 363.32 | 358.21 | 4.84 | 0.01 | 320.83 | 356.19 | 371.05 | 349.36 | 25.80 | 0.07 |
| 0.70 | 419.91 | 409.72 | 424.91 | 418.18 | 7.74 | 0.02 | 390.57 | 434.42 | 460.92 | 428.64 | 35.53 | 0.08 |
| 0.80 | 480.24 | 462.94 | 484.27 | 475.82 | 11.33 | 0.02 | 464.10 | 517.30 | 563.22 | 514.87 | 49.61 | 0.10 |

### Intra-Person and Inter-Person Variability

Intra-person variability was assessed using data from several investigators. Each investigator submitted the results from three previously run assays performed with the Pierce™ BCA Protein Assay Kit using freshly diluted BSA standards with concentrations ranging from 0-420µg/ml. Intra-person variability was higher for some investigators than others. While the standard curves all had R^2^ values > 0.99, the slopes of the curves varied from assay to assay within the data sets from two of the three investigators (Figure 3 B, C). The slopes of the curves were identical for one investigator, yielding identical standard curves and very low variability (Figure 3 A). The calculated protein concentrations from Investigator 3 were significantly different than those from Investigators 1 or 2 (Table 3), and the CV was higher for all of the samples when interpolated from Investigator 3 standard curves (0.13-0.15). The CV was very low (0.01-0.02) when Investigator 1 standard curves were used. The CV was higher for Investigator 2 than Investigator 1 but still low (0.06-0.10) when the highest and lowest hypothetical samples at the edge of the curve are excluded. It was not possible to test if the protein concentrations of actual rather than hypothetical samples run by these investigators would be the same since different samples were run in separate experiments. Since these were hypothetical samples of assigned absorbance values, the BCA developer used in these assays had no effect on the samples as they would when actual samples are used, even though background was subtracted from the absorbance of the hypothetical samples. When the same developer is used for the standard curve and the samples, the effect on absorbance of the standards and samples may be the same and the resultant protein concentrations may be more similar. This underscores the need to run a standard curve at the same time the samples are run, using the same developer.

**Table 3.**
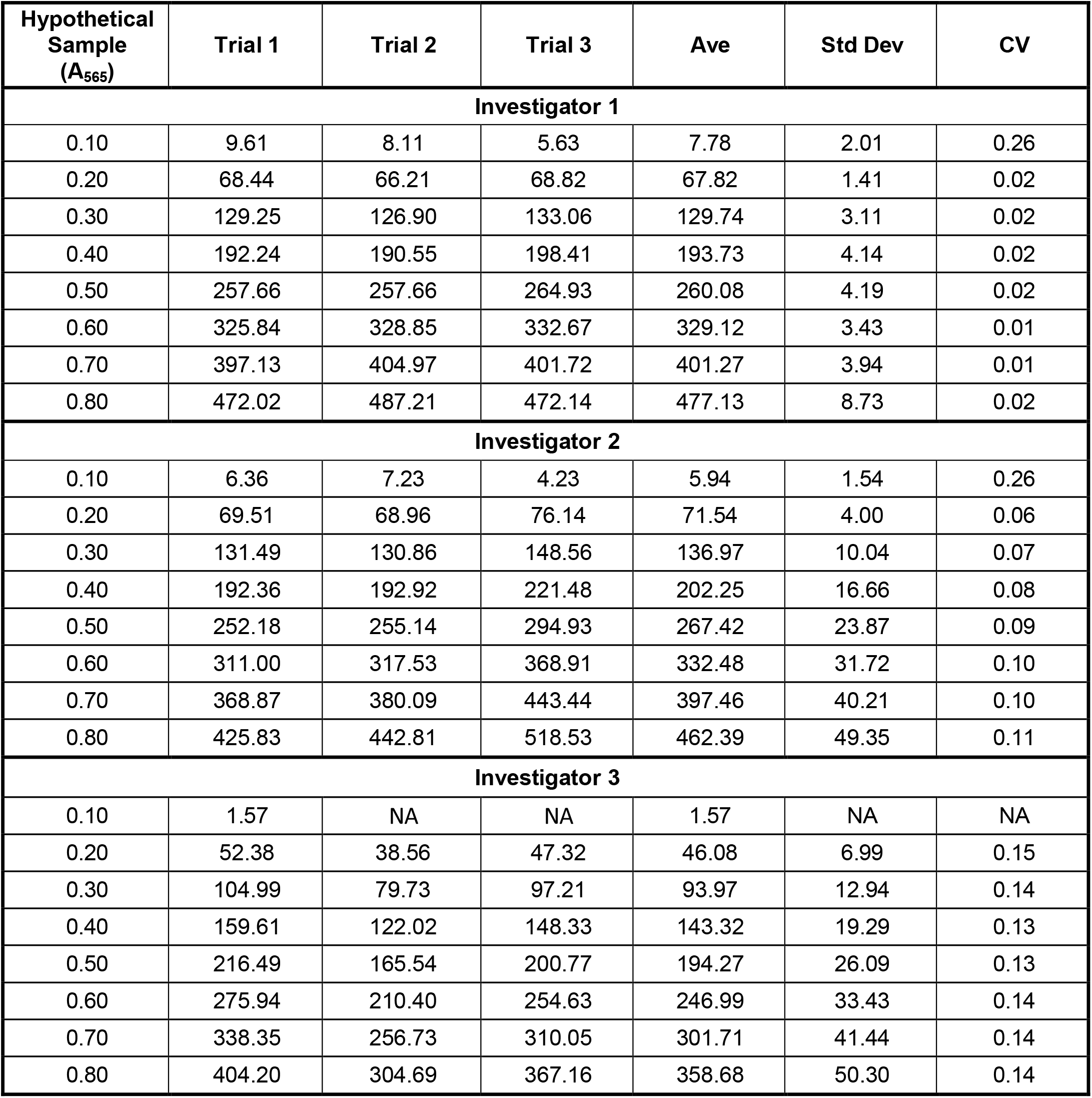
Intra-person variability was evaluated for three different investigators. Each investigator prepared three separate curves. Protein concentrations were calculated for hypothetical absorbance values using each curve.

| Hypothetical Sample (A<sub>565</sub>) | Trial 1 | Trial 2 | Trial 3 | Ave | Std Dev | CV |
| --- | --- | --- | --- | --- | --- | --- |
| **Investigator 1** |  |  |  |  |  |  |
| 0.10 | 9.61 | 8.11 | 5.63 | 7.78 | 2.01 | 0.26 |
| 0.20 | 68.44 | 66.21 | 68.82 | 67.82 | 1.41 | 0.02 |
| 0.30 | 129.25 | 126.90 | 133.06 | 129.74 | 3.11 | 0.02 |
| 0.40 | 192.24 | 190.55 | 198.41 | 193.73 | 4.14 | 0.02 |
| 0.50 | 257.66 | 257.66 | 264.93 | 260.08 | 4.19 | 0.02 |
| 0.60 | 325.84 | 328.85 | 332.67 | 329.12 | 3.43 | 0.01 |
| 0.70 | 397.13 | 404.97 | 401.72 | 401.27 | 3.94 | 0.01 |
| 0.80 | 472.02 | 487.21 | 472.14 | 477.13 | 8.73 | 0.02 |
| **Investigator 2** |  |  |  |  |  |  |
| 0.10 | 6.36 | 7.23 | 4.23 | 5.94 | 1.54 | 0.26 |
| 0.20 | 69.51 | 68.96 | 76.14 | 71.54 | 4.00 | 0.06 |
| 0.30 | 131.49 | 130.86 | 148.56 | 136.97 | 10.04 | 0.07 |
| 0.40 | 192.36 | 192.92 | 221.48 | 202.25 | 16.66 | 0.08 |
| 0.50 | 252.18 | 255.14 | 294.93 | 267.42 | 23.87 | 0.09 |
| 0.60 | 311.00 | 317.53 | 368.91 | 332.48 | 31.72 | 0.10 |
| 0.70 | 368.87 | 380.09 | 443.44 | 397.46 | 40.21 | 0.10 |
| 0.80 | 425.83 | 442.81 | 518.53 | 462.39 | 49.35 | 0.11 |
| **Investigator 3** |  |  |  |  |  |  |
| 0.10 | 1.57 | NA | NA | 1.57 | NA | NA |
| 0.20 | 52.38 | 38.56 | 47.32 | 46.08 | 6.99 | 0.15 |
| 0.30 | 104.99 | 79.73 | 97.21 | 93.97 | 12.94 | 0.14 |
| 0.40 | 159.61 | 122.02 | 148.33 | 143.32 | 19.29 | 0.13 |
| 0.50 | 216.49 | 165.54 | 200.77 | 194.27 | 26.09 | 0.13 |
| 0.60 | 275.94 | 210.40 | 254.63 | 246.99 | 33.43 | 0.14 |
| 0.70 | 338.35 | 256.73 | 310.05 | 301.71 | 41.44 | 0.14 |
| 0.80 | 404.20 | 304.69 | 367.16 | 358.68 | 50.30 | 0.14 |

Using the same standard curves from Investigators 1, 2, and 3, variability between investigators (inter-person) was also studied. The CV was higher (0.09-0.12) for inter-person variability than intra-person variability when the 0.1 hypothetical sample was excluded, with the exception of the curves from Investigator 3 which demonstrated high intra-person variability as evidenced by CV > 10% for all standard dilutions (Figure 4 and Table 4).

**Table 4.** Inter-person variability between three investigators was evaluated. Each investigator prepared three curves, and a curve was made from the average of the data for each person for comparison. Each standard was assayed in triplicate.

| Hypothetical Sample ($A_{565}$) | Investigator 1 | | | Investigator 2 | | | Investigator 3 | | | Ave | St Dev | CV |
| --- | --- | --- | --- | --- | --- | --- | --- | --- | --- | --- | --- | --- |
|  | Trial 1 | Trial 2 | Trial 3 | Trial 1 | Trial 2 | Trial 3 | Trial 1 | Trial 2 | Trial 3 |  |  |  |
| 0.10 | 9.61 | 8.11 | 5.63 | 6.36 | 7.23 | 4.23 | 1.57 | NA | NA | 6.11 | 2.64 | 0.43 |
| 0.20 | 68.44 | 66.21 | 68.82 | 69.51 | 68.96 | 76.14 | 52.38 | 38.56 | 47.32 | 61.82 | 7.23 | 0.12 |
| 0.30 | 129.25 | 126.90 | 133.06 | 131.49 | 130.86 | 148.56 | 104.99 | 79.73 | 97.21 | 120.23 | 12.84 | 0.11 |
| 0.20 | 192.24 | 190.55 | 198.41 | 192.36 | 192.92 | 221.48 | 159.61 | 122.02 | 148.33 | 179.77 | 18.08 | 0.10 |
| 0.50 | 257.66 | 257.66 | 264.93 | 252.18 | 255.14 | 294.93 | 216.49 | 165.54 | 200.77 | 240.59 | 22.99 | 0.10 |
| 0.60 | 325.84 | 328.85 | 332.67 | 311.00 | 317.53 | 368.91 | 275.94 | 210.40 | 254.63 | 302.86 | 27.79 | 0.09 |
| 0.70 | 397.13 | 404.97 | 401.72 | 368.87 | 380.09 | 443.44 | 338.35 | 256.73 | 310.05 | 366.82 | 32.86 | 0.09 |
| 0.80 | 472.02 | 487.21 | 472.14 | 425.83 | 442.81 | 518.53 | 404.20 | 304.69 | 367.16 | 432.73 | 38.80 | 0.09 |

### Intra-person and Inter-person Variability While Controlling for Assay Kit Developer

To test intra-person variability while controlling for developer variability, four sets of standards made by the same investigator (using a single BSA stock) were assayed together on one plate using the same developer. When developer variability was controlled in this way, the curves for each set of standards prepared by one investigator were aligned for protein concentrations ≤240µg/ml. At higher concentrations the slope of the curves varied slightly with increasing concentration (Figure 5). When eight absorbance values (0.1-0.8) were used as hypothetical samples and the 0.1 hypothetical sample was excluded, the CV was low (0.01-0.05) and the variability in interpolated protein concentrations was not statistically different (Table 5).

**Table 5.** Four sets of diluted standards were prepared by one investigator and were used in triplicate to create standard curves. They were assayed on the same plate to assess intra-person variability when plating and developer variability were controlled. Eight absorbance values (0.1-0.8) were used as hypothetical samples using each of the curves.

| Hypothetical Sample ($A_{565}$) | Standard Set 1 | Standard Set 2 | Standard Set 3 | Standard Set 4 | Ave | St Dev | CV |
| --- | --- | --- | --- | --- | --- | --- | --- |
| 0.10 | 4.20 | 8.77 | 3.63 | 9.84 | 6.61 | 3.15 | 0.48 |
| 0.20 | 69.14 | 68.84 | 66.72 | 69.26 | 68.49 | 1.19 | 0.02 |
| 0.30 | 134.28 | 132.00 | 132.15 | 131.01 | 132.36 | 1.38 | 0.01 |
| 0.20 | 199.60 | 198.80 | 200.19 | 195.37 | 198.49 | 2.16 | 0.01 |
| 0.50 | 265.11 | 269.94 | 271.20 | 262.72 | 267.24 | 4.00 | 0.01 |
| 0.60 | 330.82 | 346.40 | 345.60 | 333.50 | 339.08 | 8.07 | 0.02 |
| 0.70 | 396.72 | 429.59 | 423.91 | 408.32 | 414.64 | 14.95 | 0.04 |
| 0.80 | 462.82 | 521.66 | 506.84 | 487.94 | 494.82 | 25.41 | 0.05 |

To test inter-person variability while controlling for developer variability, three sets of standards were made by three different investigators at the same time. Three samples were assayed together on one plate using these standards and the same developer. Standard curves made by 2 of the 3 investigators were aligned for protein concentrations when developer variability was controlled in this way (Figure 6). When the 0.1 hypothetical sample is excluded, the CV for the samples using standards made by different investigators (0.08-0.10) was higher than the CV obtained using multiple sets of standards from one investigator (Table 6, Table 5). Inter-person variability in interpolated protein concentrations for the samples was greater than intra-person variability, although there was not a statistically significant difference between two of the investigators.

**Table 6.** Standard curves made by three investigators, plated by one person and assayed on the same plate were compared to assess inter-person variability while plating and developer variability were controlled. Each investigator made one curve in triplicate. Three samples (n=6) were assayed on the same plate and each curve was used to determine protein concentration.

| Sample<br>(A<sub>565</sub>) | Calculated protein concentration (µg/ml) |  |  |  |  |  |
| --- | --- | --- | --- | --- | --- | --- |
|  | Investigator 1<br>Standards | Investigator 2<br>Standards | Investigator 3<br>Standards | Average | St Dev | CV |
| Sample 1 | 182.11 | 195.86 | 222.25 | 200.07 | 20.40 | 0.10 |
| Sample 2 | 202.51 | 216.85 | 243.98 | 221.11 | 21.06 | 0.10 |
| Sample 3 | 260.03 | 275.46 | 302.30 | 279.26 | 21.39 | 0.08 |

## Discussion

The ability to accurately quantitate protein is critical to many areas of scientific investigation. Modern technologies for detecting protein expression have advanced, allowing smaller amounts of total protein to be used. Thus, there is an increased need to ensure that protein quantitation methods are reproducible and accurate. Commonly used methods typically involve colorimetric assays in which the sample is compared to a standard curve created from a dilution series of a known protein. However, as samples are often isolated and quantitated by different investigators in a laboratory and later used in multiple experiments, it is important to understand sources of variability in these assays and control for them whenever possible. These may include the different measuring devices used to setup the assay and intra-person and inter-person variability in the preparation of the standards and assay reagents (developers). A series of experiments was designed to determine how to best mitigate the variability inherent in these assays in a shared laboratory environment.

It is common to use the R^2^ values obtained for the standard curve as a measure of assay accuracy. The R^2^ values were high (0.982-1.0) in all of the experiments conducted for this work. However, the slope of the curves varied despite the identical standard concentrations and protocols used for making the standards. When assays were run at the same time allowing for no variability in the developer, standard curves were more closely aligned and the slopes were more similar. The fact that reproducibility is increased when developer is controlled suggests that the differences in the slope of the curves may not actually affect the interpolated protein concentrations since the developer may affect the samples and standards equally. Work done by Cortés-Ríos ^1^ showed that incubation time can affect absorbance values, thus, it is imperative that samples and standard curves be run together using the same batch of developer and the same incubation times to ensure accurate and reproducible results. They and others ^1,2^ also found that very low absorbance results (<0.07) significantly increased the relative error of the assay. They suggested that improved results are seen when the absorbance is greater than 0.1. Our data agrees with this, and in many of our experiments the CV values were higher for the 0.1 hypothetical samples. Finally, BCA assay kits typically contain a concentrated BSA solution that is diluted to create a standard curve. Cortes-Rios et al ^1^ found that the choice of protein standard could affect the results, suggesting that the use of a single standard protein such as BSA for all assays done can provide more generalizable results across different experiments and in different laboratories.

Even when the same protein is used as a standard, variability in dilution of this standard can be a source of inconsistent concentration results. This could be mitigated by using a single batch of diluted and frozen standards in an experimental series. In fact, in this work there was considerably more reproducibility in interpolated protein concentration of samples when standards from a single batch were frozen at −20°C or −80°C and thawed immediately before use. This was true for standards that were thawed and refrozen up to three times. However, care should be taken to avoid sublimation of stored frozen samples that can occur when standard microfuge tubes are used. In these experiments, the freezing tubes used had a rubber O ring which would minimize sublimation. Sealing tubes without gaskets using parafilm may achieve the same effect, but that was not tested here, nor was time in the freezer beyond more than one month.

Using a single batch of diluted and frozen standards in assays run by all investigators in a large experimental series over time may improve the comparative value of the interpolated protein concentrations for the samples. This may be especially true in laboratories where many investigators or less experienced users are running the assays. In addition, using frozen standards is more convenient and efficient than preparing the standards each time an assay is run. There may also be a reduction in waste from leftover BSA that must be discarded after the vial is opened when the entire vial of BSA is used to make the batch of standards for freezing.

These experiments suggest that the most reproducible and accurate method for determining the protein concentration of extracted samples in an experimental series over time is diluting a large batch of BSA standards, freezing them at either −20°C or −80°C, and thawing and refreezing them up to three times. In addition, the differing slopes of curves made by different investigators or the same investigator at different times despite all curves having high R^2^ values suggests that laboratories should establish not only a minimum R^2^ value for an “acceptable” assay, but also a range of acceptable slopes for their standard curves.

## Author Contributions

RS designed experiments, collected data, analyzed data, composed figures, wrote the manuscript; BR, CEB, LC, MLH collected data, revised and edited the manuscript; PS carried out statistics, reviewed and revised results; TCT designed experiments, interpreted data, reviewed, revised and edited manuscript; ACS supervised the project, designed experiments, analyzed data, composed figures, wrote and critically reviewed manuscript, funded project. All authors approved the final version of the manuscript.

## Data Availability Statement

The data sets generated for this study are available upon request to the corresponding author.

## Competing Interests

The authors declare no competing interests.

## Funding

This work was supported by grants from Students Supporting Brain Tumor Research (SSBTR to ACS), National Institutes of Health (R01NS100793 to TCT) and Phoenix Children’s Hospital Mission Support. The content is solely the responsibility of the authors and does not necessarily represent the official views of SSBTR, the National Institutes of Health or Phoenix Children’s Hospital.

## Notes

### Competing Interest Statement

The authors have declared no competing interest.

